# Dorsomedial Striatum Calcium Permeable AMPA Receptors in the Development of Aversion-Resistant Alcohol Drinking

**DOI:** 10.64898/2026.06.11.731723

**Authors:** Meredith R. Bauer, Claudia Rangel-Barajas, Yanping Zhang, Stephen L. Boehm

## Abstract

**Rationale:** Alcohol use disorder is defined by drinking alcohol despite knowledge of negative consequences, often referred to as aversion-resistant drinking (ARD). The dorsomedial (DMS) and dorsolateral striatum (DLS) are necessary for goal-directed and habitual action selection, respectively. Leading hypotheses posit that once drug use becomes compulsive, DMS dependence degrades while DLS dependence increases. This shift may be mediated by changes in synaptic weights from glutamatergic inputs.

**Objectives:** Using a combination of western-blot, micro-injections, and ex-vivo electrophysiology, we investigated the role of α-Amino-3-hydroxy-5-methyl-4-isoxazolepropionic acid receptors AMPAR, which drive glutamatergic transmission, during quinine-adulterated alcohol (QuA) drinking in the DMS and DLS across the development of ARD.

**Results:** We found that AMPAR subunit composition and function change in the DMS across the development of ARD whereby, calcium permeable (CP) - AMPARs drive behavior. Western blots revealed a negative relationship between DMS GluA1 and QuA drinking in aversion-sensitive mice and positive relationships between DMS or DLS GluA1/A2 ratios and QuA drinking in ARD mice. DMS CP-AMPAR antagonism caused an increase in QuA drinking suggesting that CP-AMPARs in the DMS prevent ARD. Ex-vivo electrophysiology of DMS spiny projection neurons (SPNs) revealed that ARD mice had a greater rectification index than aversion-sensitive mice indicating that SPNs in the DMS express more CP-AMPARs following the development of ARD.

**Conclusions:** These data provide evidence that repeated alcohol binges alter DMS CP-AMPAR activity, where initial DMS activity acts to prevent ARD but after repeated binges that result in ARD, DMS SPNs recruit CP-AMPARs.

## Introduction

Flexible adaptation of behavior in the face of negative consequences is necessary for survival. However, in drug addiction, the processes that drive adaptive behavior may be disrupted resulting in continued use of drugs despite knowledge of negative consequences. Alcohol Use Disorder (AUD) is a chronic disease that 10% of people in the U.S. suffer from and is characterized by continued use of alcohol despite negative consequences (Substance Abuse and Mental Health Services Administration; American Psychological Association 2013). Binge alcohol drinking, a risky form of alcohol misuse, is a risk factor for the development of AUD. Understanding the neural mechanisms underlying how binge alcohol drinking produces the inflexible decision making that results in AUD may aid in the prevention and treatment of this disease.

Efficient decision making relies on competition and collaboration between goal-directed and habitual brain processes. These two systems have differences in utility and cost where habit is less computationally expensive but less accurate than goal-directedness. Thus, the brain must strike a balance utilizing the less efficient but more accurate goal-directed system and the more efficient but less accurate habitual system to execute behavior. Classic experiments identified the dorsomedial striatum (DMS) as a necessary nucleus of goal-directed action selection and the dorsolateral striatum (DLS) as a necessary nucleus for habitual action selection for natural rewards (Yin et al., 2005; 2006). Further experiments demonstrated that alcohol, both acute and chronic, shifts responding for rewards to bias habitual action selection and that this is accompanied by dependence away from the DMS and toward the DLS (Dickinson et al. 2002; Corbit et al. 2012; Corbit et al. 2014; Houck and Grahame 2018). These findings from alcohol and other drugs resulted in prominent theories of addiction suggesting that continued drug use despite negative consequences is due to a disruption in the balance between habit and goal-directed processes resulting in an overreliance on habit and a loss of goal-directedness (Everitt and Robbins 2005; Everitt and Robbins 2016; Lüscher, Robbins, Everitt 2020). Thus, this continued drug use despite negative consequences is hypothesized to be due to a physiological shift away from DMS-dependent goal-directed systems and towards a strengthened engagement of DLS-dependent habit systems (Everitt and Robbins 2005; Everitt and Robbins 2016).

Continued alcohol drinking despite negative consequences, often referred to as aversion-resistant drinking (ARD), is commonly modeled by assessing the extent to which an animal will continue to drink alcohol when it is adulterated with the bitter compound quinine or despite a mild foot shock. Compelling experimental evidence demonstrating that the development of ARD is accompanied by a shift in dependence toward the DLS is demonstrated across several experiments. Pharmacological blockade of dopamine receptors in the DLS prevents shock-resistant alcohol seeking (Giuliano et al. 2019) and ablation of DLS fast-spiking interneurons reduces willingness of mice to consume quinine-adulterated alcohol (QuA) compared with controls (Patton et al. 2019). We found that intra-DLS AMPA receptor antagonism reduced front-loading of QuA (Bauer et al. 2022a) and DLS D1 receptor antagonism reduces QuA intake without altering non-adulterated alcohol drinking (Houck 2019). Together, these data line up with leading hypotheses demonstrating that the DLS is necessary for alcohol use despite negative consequences.

However, evidence demonstrating a loss of engagement in the DMS during drinking despite negative consequences is sparse. In fact, only two separate lines of experimental data support the necessary role of the DMS in ARD. First, antagonism of DMS D1 receptors reduced QuA drinking in ARD mice without altering non-adulterated drinking (Houck 2019). Genetic knock-out (KO) of Lrrk2, a negative modulator at of D1, to effectively potentiate D1 cells significantly increases cFOS and potentiated spike activity in D1 DMS compared with littermate controls (da Silva 2024). These KO mice innately drink more alcohol and more QuA than their wild type littermates, suggesting that the activity of the DMS drives alcohol drinking despite negative consequences. Together, the current state of the field surrounding continued alcohol use despite negative consequences points to involvement of both the DLS and the DMS to drive ARD.

It has long been appreciated that alcohol seeking and alcohol taking (i.e., consumption) are separate processes (Samson 1998). The prominent theories suggesting overreliance on habit are often supported by data assessing action selection (i.e., lever presses or nose pokes) where reward consumption is often not measured. Thus, experiments assessing ARD through QuA consumption are assessing a drug taking behavior, not seeking, which may rely on separate neural processes. Nevertheless, assessment of the neurobiology of continued alcohol use despite negative consequences is often conflated across drug seeking and taking. Recent review articles on drug use despite negative consequences suggest the compulsive-like drug taking may be separate from the previous identified neural circuits of action-selection and is due to extreme goal-directedness (Hogarth 2020; Lüscher et al 2020). Despite this acknowledgement, assessment of whether the neurocircuitry of compulsive drug seeking (i.e. the DMS and DLS) persists to drive compulsive drug consumption has not been assessed.

The purpose of this project was to shed light on the neurobiology of ARD by assessing the relationship between the DMS and DLS in aversion-sensitive versus ARD mice. Extending on previous models of ARD using QuA (Wolffgramm and Heyne 1991; Lesscher et al. 2010) we have found that three weeks of binge-like alcohol access under drinking-in-the-dark (DID) procedures can produce robust ARD as demonstrated by alcohol history mice drinking significantly more QuA than water history mice and alcohol history mice drinking the same levels of QuA as non-adulterated alcohol on the day prior (Bauer et al. 2021). We hypothesized that the DMS would initially act to prevent ARD but following the development of ARD the DMS would disengage and the DLS would dominate. We found AMPAR expression differed in the DMS vs the DLS in association with ARD and that the DMS initially prevents ARD but undergoes plasticity on DMS spiny projection neurons (SPNs) in association with ARD. We demonstrate the first ever evidence of CP-AMPARs causal role in ARD and reveal a dichotomous relationship between DMS CP-AMPARs and the development of ARD. Moreover, we provide evidence in support of a gain of function of both the DMS and DLS in ARD suggesting ARD may be due to extreme goal-directness.

## Methods

### Animals

Adult male and adult female C57BL/6J mice were acquired from The Jackson Laboratory (Bar Harbor, ME). Animals were individually housed in a vivarium with 12 hour:12 hour reverse light-dark cycle for 1 week prior to the start of experiments. Mice were housed in nonfiltered wired top standard shoe box mouse cages and were given food (Lab Diet 5001, Rodent Diet) and tap water ad libitum with the exception of water bottle removal during the DID sessions. Procedures were approved by the IU Indianapolis School of Science Institutional Animal Care and Use Committee and conformed to the Guide for the Care and Use of Laboratory Animals.

### Binge and Aversion-Resistant Drinking

#### Drinking-in-the-Dark (DID)

DID is a limited access model of binge-like alcohol consumption (Thiele et al. 2014). Mice received one 10 ml ball-bearing sipper tube filled with 20% v/v ethanol in tap water (Pharmco Inc, Brookfield, CT) for two hours a day, three hours into the dark cycle, in their home cage. Consumption was measured by reading the sipper tubes to the nearest 0.025 ml, and volumes were adjusted for leak based on remaining fluid measured from sipper tubes in empty cages on the same drinking rack. Body weights were taken the day prior to day 1 of DID and weekly thereafter to determine g/kg consumption. DID reliably produces blood alcohol concentrations at or above the level of a binge (i.e., ≥ 80 mg/dL, Bauer et al. 2021).

#### Aversion-Resistant Drinking (ARD)

ARD is measured by determining the level of QuA the mouse is willing to drink and comparing these levels with baseline alcohol consumption. QuA was made by adding quinine hemi-sulfate (500 μM; 0.1957 g/L) to 20% alcohol solutions. Quinine hemi-sulfate was purchased from Millipore Sigma (St Louis, MO). This concentration of QuA was chosen based on previous work in our lab finding that 500 μM is accepted in alcohol history mice to the same level as non-adulterated alcohol but is rejected in water-history animals comparatively.

### Western Blots

#### Subcellular Fractionation and Western Blots

Twenty-four hours after behavioral testing, brains were harvested and DMS and DLS regions were rapidly dissected on ice. Synaptic isolation was then performed to separate using a subcellular fractionation, followed by Western blot analyses to examine protein expression at the synapse. All fractionation and blotting procedures were carried out as previously described (Bauer et al. 2024).

### Cannula Surgery and Microinjections

#### Surgery

The DMS was targeted for bilateral canulations. Specific procedures have been described in-depth elsewhere (Bauer et al. 2022b) DMS coordinates used were M/L: ±1.25 mm, A/P: +.5 mm and D/V: −2.75 mm. Following surgery, mice received a single subcutaneous injection of 5 mg/mL carprofen (a post-operative pain treatment) at 10 mL/kg and were placed on a heating pad until recovery from anesthesia (approximately 30 min). Mice were given ad-libitum food and water and at least a week of recovery prior to DID.

#### Microinjections

Stylets were changed daily. Mice were given mock injection two days prior to microinjections immediately prior to the DID session by using microinjectors that extended .5 mm past the cannula and restraining the mouse for the 2.5 min duration of a microinjection. Mice were microinjected every other day with drug doses assigned in a Latin-square design. Mice were either injected with varying concentrations of the CP-AMPAR antagonist, NASPM (0, 20, or 40 μl/side) or M4 antagonist, Tropicamide, (0, 1, or 3 μl/side) immediately prior to DID.

#### Placement Verification

Following the completion of the behavioral data collection, brains were taken for verification of cannula placement. Mice were cervically dislocated; brains were extracted and flash-frozen in 2-methylbutane at −20 to −40°C. Brains were sectioned at 40 μm, slices were stained with cresyl violet and placements were determined by a blinded experimenter.

### Whole-Cell Voltage-Clamp Recordings

#### Brain slice preparation

Mice were decapitated under deep isoflurane anesthesia, and the brain was immediately removed and placed in oxygenated (95% O2/5% CO2), ice-cold cutting solution consisting of the following (in mM): 194 sucrose, 30 NaCl, 4.5 KCl, 1.2 NaH2PO4, 26 NaHCO3, 1 MgCl2, and 10 glucose, at pH 7.4; 310 mOsm. Brain slices were cut using a vibrating-blade microtome (model VT 1200S, Leica Biosystems). Immediately after cutting, slices were kept at room temperature for a resting period of at least 1 h in artificial CSF (aCSF) solution containing the following (in mM): 124 NaCl, 2.5 KCl, 25 NaHCO3, 1.25 NaH2PO4, 2 CaCl2, 1 MgCl2, and 25 glucose, at pH 7.4; 290 mOsm.

#### Electrophysiological recordings

Whole-cell voltage-clamp recordings were performed as previously reported (Rangel-Barajas et al. 2021). After a resting period following slice preparation, individual coronal slices containing the DMS were transferred to a recording chamber mounted on an upright microscope and continuously perfused with oxygenated artificial cerebrospinal fluid (aCSF, 2 mL/min) at room temperature (∼22–24°C). Patch pipettes (resistance 3–6 MΩ) were fabricated from borosilicate glass capillaries (1.5 mm outer diameter, 1.12 mm inner diameter) using a horizontal puller (P-1000, Sutter Instruments, Novato, CA) and filled with a CsMeSO_3_-based internal solution containing (in mM): 120 CsMeSO3, 5 NaCl, 10 HEPES, 1.1 EGTA, 0.3 Na-GTP, and 4 Mg-ATP (pH 7.4, 290 mOsm). All reported membrane potentials were corrected for the liquid junction potential (10 mV) calculated using the pClamp junction potential calculator. Cells were visualized with a Nikon Eclipse FN-1 microscope equipped with a 40x water-immersion objective and an ORCA-FLASH sCMOS camera (Hamamatsu) using differential interference contrast imaging. Recordings were obtained using a MultiClamp 700B amplifier and Digidata 1550B digitizer (Molecular Devices, Sunnyvale, CA), filtered at 2 kHz, digitized at 10 kHz, and acquired with Clampex 11 software (Molecular Devices, Sunnyvale, CA). For all recordings, series resistance was 10–20 MΩ and was left uncompensated, recordings were excluded if series resistance change by >20%.

SPNs in the DMS were identified based on visual examination of morphological (oval soma of ∼10 µm in diameter) and electrophysiological characteristics (i.e., capacitance and membrane resistance) parameters (Rangel-Barajas 2021). Basic membrane properties were determined using a 5 mV depolarizing voltage step with the membrane test function in pClamp. Neurons were allowed to equilibrate in whole-cell configuration for 5-10 minutes prior to recording to allow for intracellular equilibration of internal solution. Evoked excitatory postsynaptic currents (eEPSCs) were recorded in aCSF containing bicuculline (10 µM) and the NMDA receptor antagonist AP-5 (50 µM) to isolate AMPA receptor-mediated responses. Synaptic stimulation was delivered via a bipolar electrode positioned ∼200–300 µm from the recording site in the corpus callosum ipsilateral to the recorded neuron. Stimulus intensity (100–320 µA) was adjusted to evoke EPSCs of 150–350 pA across at least six consecutive pulses, with <10% variability between pulses. NASPM-sensitive currents were quantified as the percent reduction from baseline. CP-AMPAR-mediated currents were pharmacologically isolated by bath application of 1-naphthyl acetyl spermine (NASPM, 100 µM) for 20 minutes, during which eEPSCs were continuously recorded.

Synaptic CP-AMPA were identified based on inward rectification of AMPA-mediated EPSCs at positive membrane potentials. To assess rectification, eEPSCs were recorded at holding potentials ranging from –80 to +60 mV in 20 mV increments. The CsMeSO3 internal solution included 100 µM spermine to maintain intracellular polyamine block. Current-voltage (I-V) relationships were generated from peak EPSC amplitudes at each holding potential, and the inward rectification index was calculated as the ratio of the current at –60 mV to that at +40 mV (I_–60_/I_+40_ mV).

### Experimental design and statistical analyses

All data were tested for normality (Shapiro-Wilk test) and homogeneity of variance (Levene’s test) and analyses were adjusted based on the results of these tests. Statistical tests used were ANOVA, t-tests, correlation (Pearson and Spearman Rank), and multiple linear regression. Bonferroni corrections on the p-value were used for correction in cases of multiple comparisons. Sex was used as a factor across all analyses and collapsed across if there were no significant differences. Differences were considered significant at p < 0.05 and magnitude of effect is reported using generalized eta squared (η^2^g) and interpreted using Cohen’s d. Figures were made using MATLAB and Adobe Illustrator 2025. The data were analyzed using R Studio (www.r-project.org).

## Results

### Post-Synaptic Density Enriched AMPA Receptor Protein Expression in the DMS and DLS in Aversion Resistant and Aversion Sensitive Mice

Previous unpublished data from our lab found that QuA intake was not related to AMPAR expression in either DMS or DLS whole-tissue sections in ARD or aversion-sensitive mice (data not shown). We hypothesized that any AMPAR changes that might be occurring to drive behavior would be happening at the post-synaptic density (PSD) where AMPARs are modulated during plasticity. Thus, we assessed PSD-enriched AMPAR expression in the DMS and DLS of ARD and aversion-sensitive mice. Male and female C57BL/6J mice drank either water or 20% alcohol under the DID procedure for 21 days (n = 6-8/group). On day 22, all mice were given QuA to test for ARD. Baseline alcohol drinking is shown in figure 1A. We ran separate RM ANOVAs of sex x day for each drinking-history group as the groups received different solutions across the 21 days of baseline drinking. ANOVA of sex and day in the water history mice revealed a main effect of day, F(1,14) = 7, p = 0.01, η²g = 0.12, but no significant effect of sex or significant interaction of day and sex (p’s > 0.05), Figure 1A. ANOVA of sex and day in the alcohol history mice revealed a main effect of day, F(1,14) = 12.60, p = 0.003, η²g = 0.13, but no significant effect of sex or significant interaction of day and sex (p’s > 0.05), Figure 1B. We next assessed the effect of an alcohol vs water drinking history on ARD drinking (Figure 1C). RM two-way ANOVA of sex and drinking history on QuA drinking revealed a main effect of drinking history, F(1,28) = 4.67, p = 0.04, η²g = 0.14, and no significant effect of sex or interaction of sex and drinking history (p’s < 0.05). Next, we assessed the extent of the ARD in the alcohol history mice by comparing consumption of alcohol on the day prior to QuA access with QuA consumption. RM two-way ANOVA of sex and solution revealed no significant effect of sex, solution, or interaction of sex and solution (p’s > 0.05), Figure 1D. To assess the effect that ARD has on AMPAR plasticity in the dorsal striatum, we assessed PSD-enriched GluA1 and GluA2 in the DMS and DLS of ARD and aversion-sensitive mice (Figure 1D). ANOVA of sex and drinking history for DMS GluA1, DMS GluA2, DLS GluA1, or DLS GluA2 revealed no significant effect (p’s > 0.05), Figure 1E. However, given the heterogeneity of the drinking and AMPAR expression, we sought to identify whether QuA consumption across each drinking history group predicted AMPAR expression. Bonferroni corrected correlations found a trend toward a moderate negative relationship between DMS GluA1 and QuA consumption in aversion-sensitive mice, (r_S_ = -.59, n = 16, p = 0.06), Figure 1F. This relationship suggests that DMS GluA1 AMPAR expression may be protective against QuA consumption in aversion-sensitive mice. There were no significant relationships for DMS GluA2, DLS GluA1, or DLS GluA2 (Figures 1G-I). Finally, we assessed GluA1/A2 ratios in the DMS and DLS of ARD and aversion-sensitive mice. ANOVA of sex and drinking history for DMS GluA1/A2 ratio and DLS GluA1/A2 ratio revealed no significant effect (p’s > 0.05), Figure 1J. Bonferroni corrected correlation of DMS GluA1/A2 ratio and QuA intake revealed a strong positive relationship in female ARD mice, (r_S_ = 0.9, n = 8, p = 0.02), Figure 1K. Bonferroni corrected correlation of DLS GluA1/A2 ratio and QuA intake also revealed a strong positive relationship in female ARD mice, (r_S_ = 0.9, n = 8, p = 0.02), Figure 1L. No such relationships were found in male ARD mice. These relationships suggests that female ARD mice have a greater overall GluA1/A2 ratio, which would bias toward CP-AMPARs, in both the DMS and DLS which may be causally related to the extent that mice will drink QuA. This relationship is specific to QuA as baseline alcohol consumption (i.e. the day prior to QuA testing) did not correlate with DMS GluA1, A2, or A1/A2 ratios or DLS GluA1, A2, or A1/A2 ratios in the alcohol history mice (data not shown). Overarchingly, these data suggest a relationship between CP-AMPARs and ARD, in dichotomous ways by drinking history or by sex.

**Fig 1.**
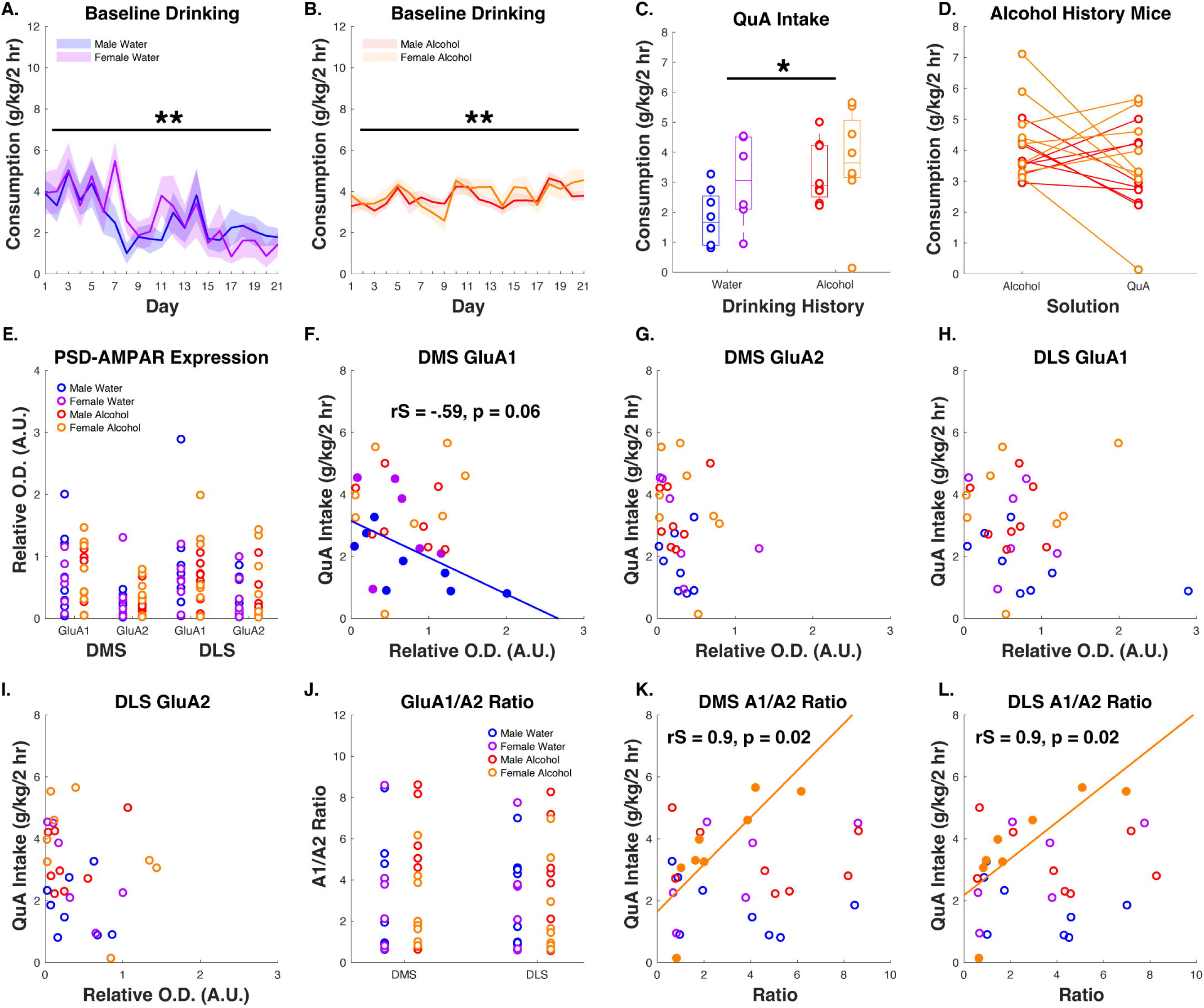
PSD-Enriched AMPAR Expression Predicts QuA Consumption. Male and female C57BL/6J mice drank either water or 20% alcohol under the DID procedure for 21 days. On day 22 all mice were given QuA to determine the extent of ARD. DMS and DLS PSD-enriched sections were assessed with Western-Blot to quantify GluA1 and GluA2 AMPA receptor subunits. **A.** Average daily water consumption in aversion-sensitive mice across the 21 days. RM two-way ANOVA revealed a main effect of day (**p < 0.01). **B.** Average daily alcohol consumption in the ARD mice across the 21 days. RM two-way ANOVA of day and sex revealed a main effect of day (**p < 0.01). **C.** QuA consumption in ARD and aversion-sensitive mice. Two-way ANOVA of sex and drinking history revealed a main effect of drinking history (*p < 0.05). **D.** Consumption in alcohol history mice comparing QuA intake with alcohol intake on the day prior to testing. RM two-way ANOVA of sex and solution revealed a trend toward a main effect of sex (p = 0.06). **E.** GluA1 and GluA2 AMPA receptor values for DMS and DLS sections in aversion-sensitive and ARD mice. ANOVAs in each brain region reveal no significant effects of sex, history, or protein for DMS or DLS sections. **F.** Bonferroni corrected correlations reveal DMS GluA1 trends toward a negative relationship with QuA intake (p = 0.06). **G-I.** DMS GluA2, DLS GluA1, and DLS GluA2 reveal no significant relationships with QuA intake. **J.** GluA1/A2 ratios were not different across sex or drinking history. **K-L.** DMS and DLS A1/A2 ratios positively and significantly predicted QuA intake in female ARD mice (p’s < 0.05). Data are displayed as individual data points or as mean ± SEM

### DMS CP-AMPAR Antagonism Causes ARD

Next, we sought to causally test whether the PSD-enriched DMS GluA1 negative relationship with QuA intake in aversion-sensitive mice is preventing ARD. Relative increases in GluA1 suggest the presence of CP-AMPA receptors. CP-AMPA receptors are most commonly GluA2 lacking GluA1 homologs and are implicated in drug addiction (Cull-Candy and Farrant 2021). Further, overarching theories of addiction suggest that the DMS is required for goal-directed, non-compulsive behavior. With this in mind, it would make sense that CP-AMPA receptors could result in greater excitability within the DMS during QuA drinking, causing goal-directed, aversion-sensitive. Thus, we hypothesized that the negative correlation between GluA1 PSD AMPAR expression and QuA drinking is due to the presence of CP-AMPARs. Male and female (n = 6/sex) mice drank water in the DID paradigm for 21 days, consistent with the drinking history in our Western-Blot data (Figure 1). On days 23, 25, and 27 mice were bilaterally infused with the CP-AMPAR antagonist, NASPM (0, 20, or 40 ug/side), into the DMS, in a within-subjects Latin-square design immediately before QuA access. Because alcohol naïve mice lack tolerance to alcohol, we wanted to be sure to capture any increases in alcohol drinking that might occur early in a drinking session as a result of the NASPM and prior to when the pharmacological action of alcohol took place potentially causing a sedative effect. Thus, bottles were read at 15-minutes in addition to being read at 2 hours. RM two-way ANOVA of sex and NASPM concentration at 15-minutes revealed a main effect of NASPM, F(1,10) = 7.01, p = 0.02, η²g = .22, but no significant effect of sex or interaction of sex and NASPM concentration (p’s > 0.05), Figure 2B. Post-hoc Dunn’s test with FDR corrections revealed a significant difference between 0 NASPM and 20 (p = 0.007) and trended toward significant difference between 0 and 40 ug/side NASPM (p = 0.059). Importantly, there was not a significant effect of order of infusion/QuA presentation which suggests that our identified effect of NASPM on QuA intake is not due to novelty of the QuA solution (p > 0.05). RM two-way ANOVA of sex and concentration of NASPM at 2-hours revealed no significant effects, Figure 2C. RM two-way ANOVA assessing the proportion of intake in 15 minutes of the 2-hour session across sex and NASPM concentration found a significant effect of NASPM concentration F(1,10) = 5.09, p = 0.047, η²g = .13, Figure 2D. Post-hoc Dunnet’s test revealed that 0 and 20 ug/side NASPM significantly differed from one another (p = 0.007), but 0 vs 40 did not (p > 0.05). These data suggest that NASPM increased QuA intake and these NASPM-induced increases were responsible for a majority of the QuA intake over the session. BACs were taken immediately following the DID session on the last infusion day (Figure 2E). Figure 2F displays a bubble chart of BACs, QuA consumption at 2 hours, and QuA consumption at 15 minutes. Bubble size display BAC, where larger bubbles indicate a higher BAC and smaller bubbles indicate a lower BACs. Pearson’s correlation between BAC and 2-hour QuA intake revealed a significant positive correlation, r(10) = .71, p = 0.009 (Figure 2F). There was no significant relationship between BAC and 15-minute QuA intake (Figure 2F) which makes sense as bloods were taken only after the 2-hour timepoint. We also assessed ambulatory locomotor activity in the home cage during the DID sessions on infusion days. RM two-way ANOVA of sex and NASPM concentration at the fifteen-minute time point revealed a main effect of drug concentration, F(1,10) = 5.27, p = 0.045, η²g = .12, Figure 2G. Post-hoc Dunn’s test revealed that there is a significant difference between the saline infusion and the 20 ug/side concentration of NASPM (p = 0.002), but not 40. Mixed effects ANOVA of locomotor activity across time during the entire two-hour session revealed no significant effect of drug concentration across time, however drug concentration and time bin significantly interacted, F(1,781) = 4.89, p = 0.027, η²= .006, Figure 2G. Assessment of total locomotor activity (area under the curve) of the two-hour session revealed a main effect of NASPM concentration, F(1,10) = 7.8, p = 0.019, η²g = 0.2, Figure 2H. Post-hoc Dunnet’s test revealed that neither 20 (p = 0.054) or 40 μg/μl NASPM (p > 0.05) differed from 0. To confirm the specificity of intra-DMS NASPM on ARD, we tested whether intra-DMS NASPM at our most effective concentration (20 μg/side) would alter non-specific consummatory behavior by measuring changes in water consumption. We found that intra-DMS NASPM did not alter water drinking or ambient locomotor activity at 15-minutes or 2 hours (p’s > 0.05), Figures 2I-L. The lack of locomotor effect of NASPM in the water condition suggests that the intra-DMS increases in locomotor activity during QuA drinking are the byproduct of ethanol induced stimulation via rapid increases in QuA consumption, and not due to CP-AMPAR antagonism. Taken together, these results demonstrate that 1) DMS CP-AMPA receptors prevent ARD, 2) this effect is unrelated to novelty of stimulus, and 3) this effect is somewhat selective as water consumption was unaffected by intra-DMS NASPM. These data also show compelling evidence that CP-AMPA receptors within the DMS gate ARD and provide support for our initial hypotheses that the DMS is engaged during early, non-ARD drinking, acting to prevent ARD.

**Fig 2.**
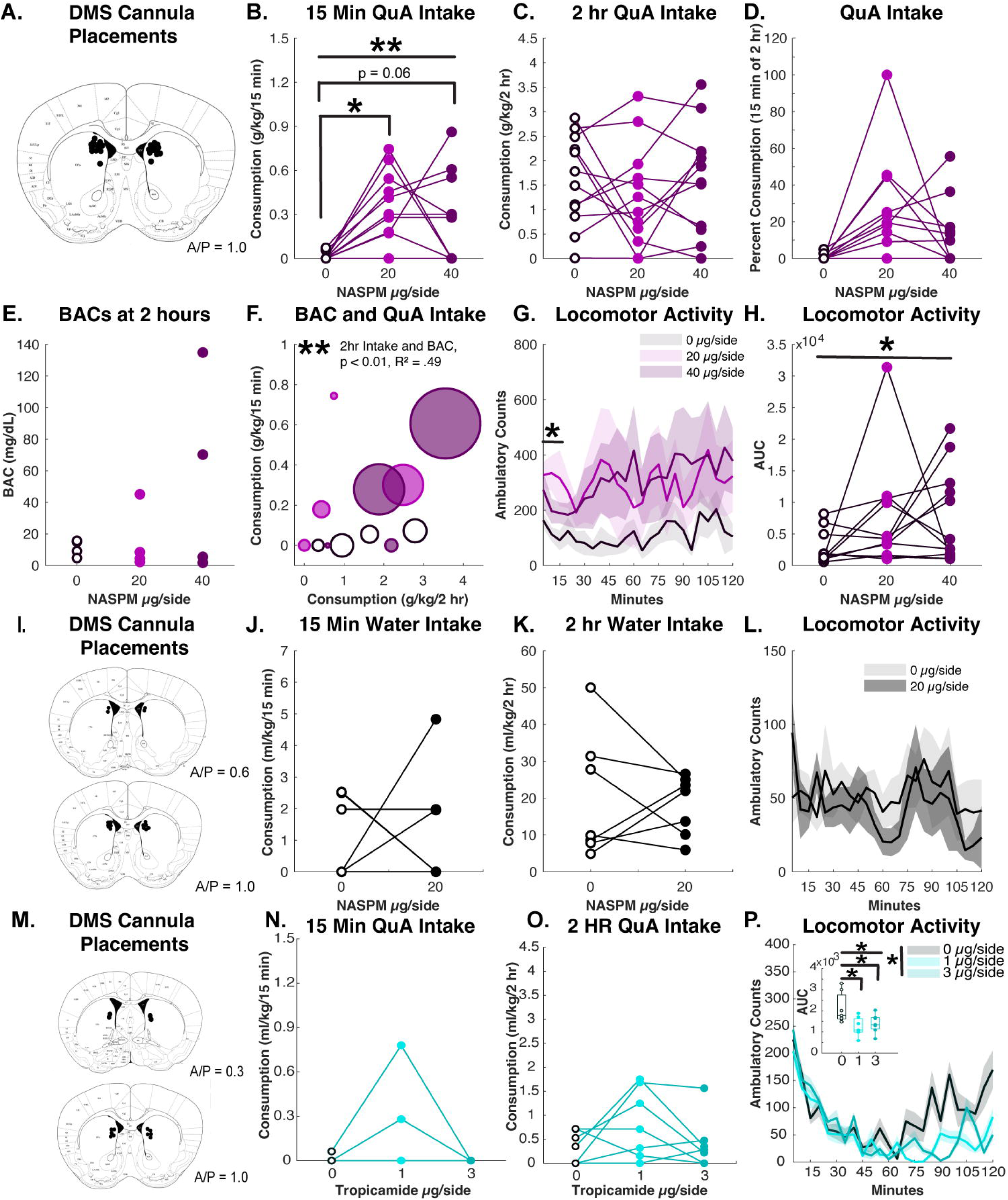
DMS CP-AMPARs Selectively Prevent ARD. **A.** DMS cannula placements for the NASPM experiment. **B.** 15-minute QuA intake across NASPM concentrations. RM one-way ANOVA revealed a main effect of concentration (**p < 0.01). Post hoc paired t-test revealed a significant difference between 0 and 20 (* p < 0.05). **C.** 2-hour QuA intake across NASPM concentrations. RM one-way ANOVA revealed no significant differences. **D.** NASPM mice consumed a large proportion of their total QuA intake in 15 minutes (0 vs 20, * p < 0.05). **E.** BACs taken immediately after the final DID session**. F.** BACs positively and significantly predicted 2-hour intake (**p < 0.01). **G-H.** NASPM significantly increased locomotor activity at both 15 minutes (*p < 0.05) and 2-hours (*p < 0.05). **I.** DMS cannula placements for NASPM water control experiments. **J-L**. NASPM did not alter water consumption or locomotor activity during the DID session. **M.** DMS cannula placements for the tropicamide experiment. N-**O.** Tropicamide did not alter QuA consumption. **P.** Intra-DMS tropicamide reduced locomotor activity (*p < 0.05). Data are displayed as individual data points or as mean ± SEM.

Next, to confirm the specificity of CP-AMPARs for this effect we antagonized DMS M4 receptors and tested ARD. M4 receptors are highly expressed in the DMS both on D1 SPNs, cholinergic interneurons, and presynaptically from cortical and thalamic inputs (Moehle and Conn 2019). Intra-DMS tropicamide (M4 antagonist) did not alter QuA consumption at 15-minutes or at two hours (p’s > 0.05), Figures 2N-O. However, ANOVA revealed a significant interaction of time and tropicamide concentration on locomotor activity, F(1,781) = 4.89, p = 0.027, η²g = 0.006, Figure 2P. RM one-way ANOVA on the area under the curve revealed a main effect of tropicamide, F(1,10) = 5.44, p = 0.04, η²g = 0.35. Post hoc Bonferroni corrected Dunnett’s paired t-test revealed a significant difference between 0 and 1 (p < 0.001) and 0 and 3 (p < 0.001) where tropicamide decreases locomotor activity, inset of Figure 2P. These data increase confidence in the specificity of DMS CP-AMPARs preventing ARD. However, it is possible that DMS tropicamide has a multiplexing effect on both locomotion (inhibiting locomotion) and QuA (increasing drinking), with the later effect being masked by a broad reduction in movement.

### Relationship Between CP-AMPARs, DMS SPNs, and QuA Drinking

Because SPNs are the primary cell type in the DMS and are affected by alcohol exposure (Rangel-Barajas et al. 2021) we sought to assess whether SPNs neurons in the DMS are expressing CP-AMPARs and whether ARD and aversion sensitive alter SPN CP-AMPAR activity. To do this, we performed whole-cell voltage clamp electrophysiology on SPNs in the DMS in aversion sensitive and ARD mice 24 hours after QuA access (n = 4 mice/sex/drinking history). Mice drank either water (aversion sensitive) or alcohol (ARD group) for three weeks. There was no significant effect of sex or day on consumption in either treatment group (Figure 3A). On day 22 mice were tested for ARD which was determined by measuring the extent to which mice would consume QuA. RM two-way ANOVA of sex and drinking history revealed that alcohol history mice drank significantly more QuA than water history mice, F(1,12) = 9.71, p = 0.009, η²g = 0.07, Figure 3B. We then assessed alcohol vs QuA consumption in the alcohol history mice only. RM ANOVA of sex and solution in the alcohol history mice did not reveal any significant effects (all p’s > 0.05), Figure 3C. 24-hours after QuA access, ex-vivo electrophysiology was performed on SPNs. Representative traces are shown in Figure 3D. RM ANOVA on the percent change of eEPSC amplitude of sex, time (pre vs post NASPM) and drinking history revealed a main effect of drinking history, F(1, 43) = 49.30, p < 0.001, η²g = 0.31, time, F(1, 43) = 41.31, p < 0.001, η²g = 0.37, interaction of sex and drinking history, F(1, 43) = 6.9, p = 0.01, η²g = 0.06, and an interaction of drinking history and time, F(1, 43) = 24.58, p < 0.001, η²g = 0.26, Figure 3E. We then ran separate RM ANOVAs for each drinking history. In the alcohol history mice, RM ANOVA of sex and time revealed a main effect of time, F(1,16) = 191.00, p < 0.0001, η²g = 0.86, sex, F(1,16) = 16.92, p < 0.0001, η²g = 0.33, and an interaction of sex and time, F(1, 16) = 15.57, p < 0.0001, η²g = 0.34. Post-hoc Dunn’s test in males (p < 0.0001) and females (p < 0.001) revealed a significant difference pre vs post-NASPM. In the water history mice, RM ANOVA of sex and time revealed no significant effects. To explicitly test the effect of drinking history and sex on EPSCs, we assessed EPSC as a percent change from baseline for each animal from timepoint 1 to timepoint 2 (see figure 3E), Figure 3F. Two-way ANOVA of sex and treatment revealed a main effect of sex, F(1,20) = 116.04, p < 0.0001, η²g = 0.85, drinking history, F(1, 20) = 1625.18, p < 0.0001, η²g = 0.99, and a significant interaction of sex and drinking history, F(1, 20) = 148.72, p < 0.0001, η²g = 0.88. Post-hoc Dunn’s tests revealed a sex difference in alcohol history mice (p < 0.01), a drinking history difference between male mice (p < 0.039), and a drinking history difference between female mice (p < 0.0039), Figure 3F. These results demonstrate that the alcohol drinking history mice had significantly greater sensitivity to NASPM suggesting that the alcohol drinking history biases DMS SPNs towards expressing greater levels of CP-AMPARs than the water history animals. These data corroborate the western-blot data in Figure 1 showing a positive relationship between GluA1/A2 ratios (likely CP-AMPARs) in female ARD mice and suggest a potential mechanism between DMS ARD and CP-AMPARs. However, we found that male ARD mice have greater SPN NASPM sensitively than female mice and male mice did not show any relationship with AMPAR expression. Additionally, these data complicate interpretation of our findings in Figures 1 and 2 showing CP-AMPARs may prevent ARD in aversion-sensitive mice.

**Fig 3.**
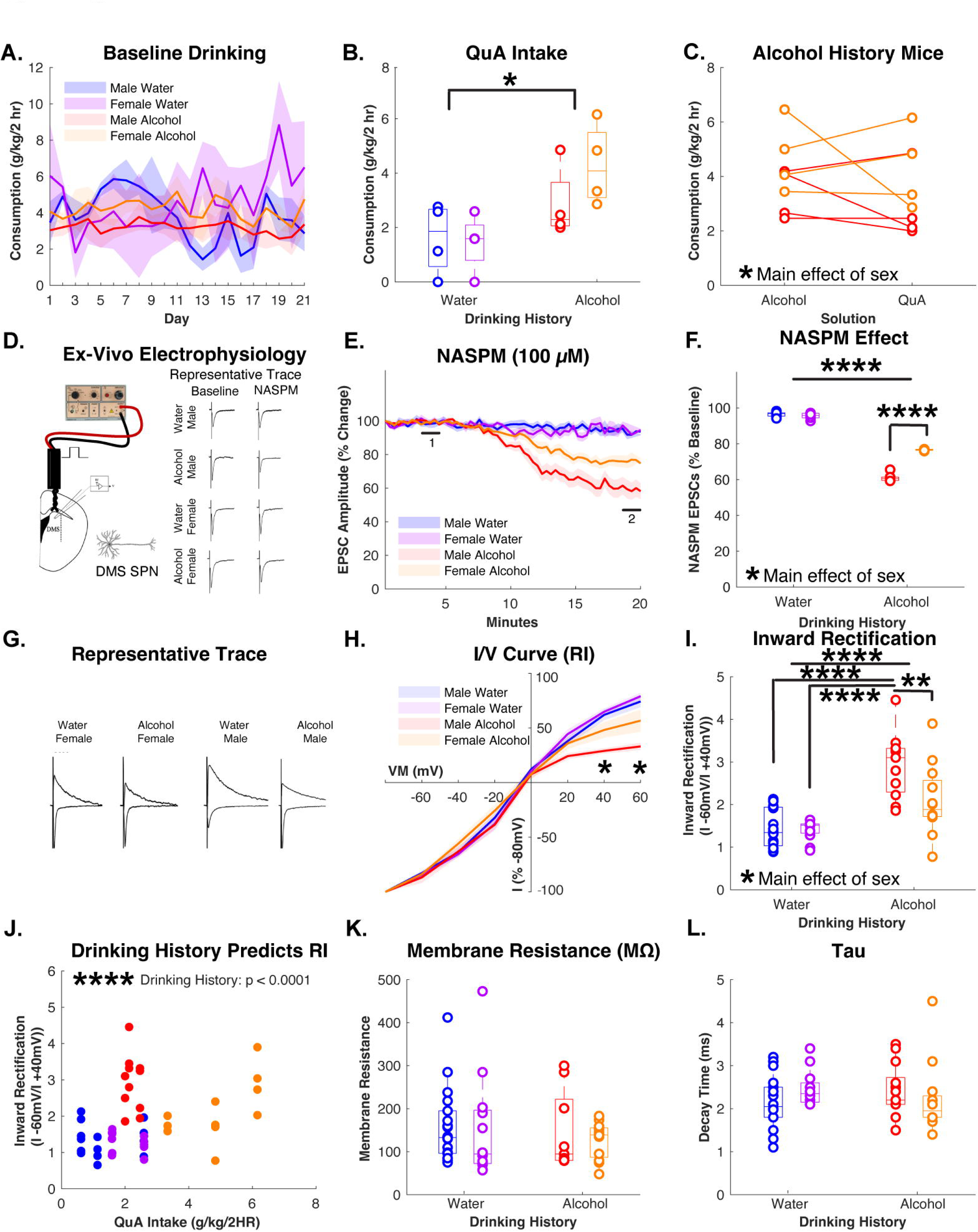
DMS SPNs are More Sensitive to NASPM in ARD Compared with Aversion-Sensitive Mice. **A.** Male and female alcohol and water history mice drank for three weeks. **B.** on day 22 both groups were given QuA. RM ANOVA revealed a main effect of drinking history (p* < 0.05). **C.** Alcohol history animals consumed the same amount of QuA as nonadulterated alcohol at baseline. Males drank less on average than females (*p < 0.05). **D.** Schematic and representative traces of our ex-vivo electrophysiology recordings. **E.** EPSC’s demonstrating a change from baseline for both drinking history groups following NASPM with greater reductions in alcohol history mice. **F.** NASPM effect as displayed as % baseline showing that alcohol history animals’ SPNs are more sensitive to NASPM than water history (***p < 0.001) and that male alcohol history mice are more sensitive to NASPM compared to female rats (****p < 0.0001). **G.** Representative traces of evoked responses. **H.** I/V curve for male and female SPNs for water and alcohol history mice. Alcohol history rats had greater RI than water history overall (p < 0.0001) and males alcohol history had greater RI than females at 40 mV (**p < 0.01) and 60 mV (**p < 0.01). **I.** Inward rectification was greater in alcohol history rats compared with water history (****p < 0.0001) and greater in males relative to females (*p < 0.05). Male alcohol history mice had greater RI than females (*p < 0.05). Males had a higher rectification index than females in both the alcohol history (****p < 0.0001) and water history (*** p < 0.001). **J.** Multiple regression analyses demonstrate drinking history significantly predicts RI (***p < 0.0001). **K-L.** There were no significant effects for membrane resistance or Tau. Data are displayed as individual data points or as mean ± SEM.

Next, we assessed the rectification index (RI) by recording evoked EPCSs at multiple holding potentials and constructing current-voltages (I/V) relationships. Representative traces are shown in figure 3G. Mixed effects ANOVA of sex, drinking history, and holding potential revealed a main effect of holding potential, F(1,350) = 5162.15, p < 0.0001, η²g = 0.937, sex, F(1,52.43) = 6.75 p = 0.01, η²g = 0.11, drinking history, F(1,52.43) = 20.42 p < 0.0001, η² = 0.28, and interaction of holding potential and sex, F(1,350) =7.73 p = 0.0057, η²g = 0.022, holding potential and drinking history, F(1,350) =43.64 p = 0, η²g = 0.11, and sex and drinking history, F(1,52.43) = 4.97 p = 0.03, η²g = 0.087, Figure 3H. To probe these interactions, we assessed the I/V curve separately by drinking history and then by sex. First, mixed effects ANOVA of sex and holding potential in the alcohol history mice showed a main effect of holding potential, F(1,161) =1569.97 p = 0, η²g = 0.907, sex, F(1,24.45) =11.05 p = 0.0028, η² = 0.31, and an interaction of holding potential and sex, F(1,161) = 6.33 p = 0.013, η²g = 0.038. Post-hoc Dunn’s test between male and female alcohol history animals at each holding potential revealed a significant difference at +40 (p = 0.01) and +60 mV (p < 0.01). Next, we assessed the effect of sex and holding potential in the water history animals only. Mixed effects ANOVA of holding potential and sex in the water history animals found a significant effect of holding potential, F(1,189) = 4289.84 p < 0.0001, η²g = 0.958. Finally, we compared holding potential and drinking history by sex. In males, a mixed-effects ANOVA of holding potential and drinking history revealed a main effect of holding potential, F(1,189) = 2752.78 p = 0, η² = 0.936, drinking history, F(1,28.29) =25.75 p = 0, η² = 0.48, and a significant interaction of holding potential and drinking history, F(1,189) = 36.64 p = 0, η²g = 0.16. Post-hoc Dunn’s test comparing drinking history for males at each holding potential revealed a significant difference between water and alcohol history groups at 20, 40, and 60 mV (all p’s < 0.05). Finally, we tested whether there was a relationship between holding potential and drinking history in female mice. Mixed-effects ANOVA revealed a main effect of holding potential, F(1,161) =2407.13 p = 0, η² = 0.937 and a significant interaction of holding potential and drinking history, F(1,161) =11.88 p < 0.0001, η² = 0.07. Post-hoc Dunn’s test at each holding potential in females revealed a significant difference between water and alcohol history mice at 40 and 60 mV (all p’s < 0.01), Figure 3H. We next assessed RI. Two-way ANOVA of sex and drinking history revealed a main effect of drinking history on RI, F(1, 47) = 24.60, p < 0.0001, η²g = 0.34, Figure 3I. Post-hoc Bonferroni corrected t-test in the alcohol history mice revealed a significant difference between males and females (p < 0.05). Separate post-hoc Wilcox rank sum test revealed a significant difference between male alcohol and water history animals (p < 0.0001) and female alcohol and water history animals (p < 0.01), 3I. Together, these data provide further evidence that the alcohol drinking history causes greater rectification index, and thus greater CP-AMPAR insertion in DMS SPNs compared with water drinking mice, and this effect was greater in males than in females.

Next, multiple regression analyses was used to test if sex, drinking history, or QuA intake significantly predicted RI in an attempt to better understand how our variables are related to DMS SPN calcium permeability. Multiple linear regression was used to test if sex, drinking history, and QuA intake predicted DMS SPN RI. The results indicated that drinking history significantly predicted 52% of the variance (R2 = 0.53, F(1,45) = 20.33, p < 0.0001), while sex and QuA intake were not significant predictors (p’s > 0.05); Figure 3J. There were no significant differences in membrane resistance (Figure 3K) or t (Figure 3L). Together, these data indicate that history of binge alcohol drinking relates to increased RI independent of sex or QuA intake. Importantly, the lack of significance of QuA intake predicting RI suggests that the DMS SPN RI may be related to alcohol consumption broadly and not necessary to ARD/QuA consumption. These results could suggest that cell types outside of SPNs are related to driving ARD.

### DMS D1/D2 Receptor Ratios Predict Alcohol Consumption, but not QuA Intake

Since we did not isolate primarily D1 or D2 expressing SPNs in the slice electrophysiology experiment, we sought to gain insight into whether relative D1 and D2 expression is related to ARD. We ran western blots on whole-tissue sections from the DMS and DLS from a drinking experiment previously published where we demonstrated aversion-sensitive and ARD (Bauer et al. 2021). Male and female C57BL/6J mice drank either water or 20% alcohol under the DID procedure for 21 days. On day 22, all mice except for the water control group were given QuA to test for ARD. We ran separate RM ANOVAs of sex x day for each drinking-history group as the groups received different solutions across the 21 days of baseline drinking. ANOVA of sex and day in the water history mice revealed no significant effects of sex, day, or significant interaction of day and sex (p’s > 0.05), Figure 4A. Mixed-effects ANOVA of sex and day in the alcohol history mice revealed a main effect of day, F(1,340) = 45.87, p < 0.001, η²g = 0.12, and a main effect of sex, F(1, 57) = 4.42, p = 0.04, η²g = 0.07, Figure 4B.

**Fig. 4.**
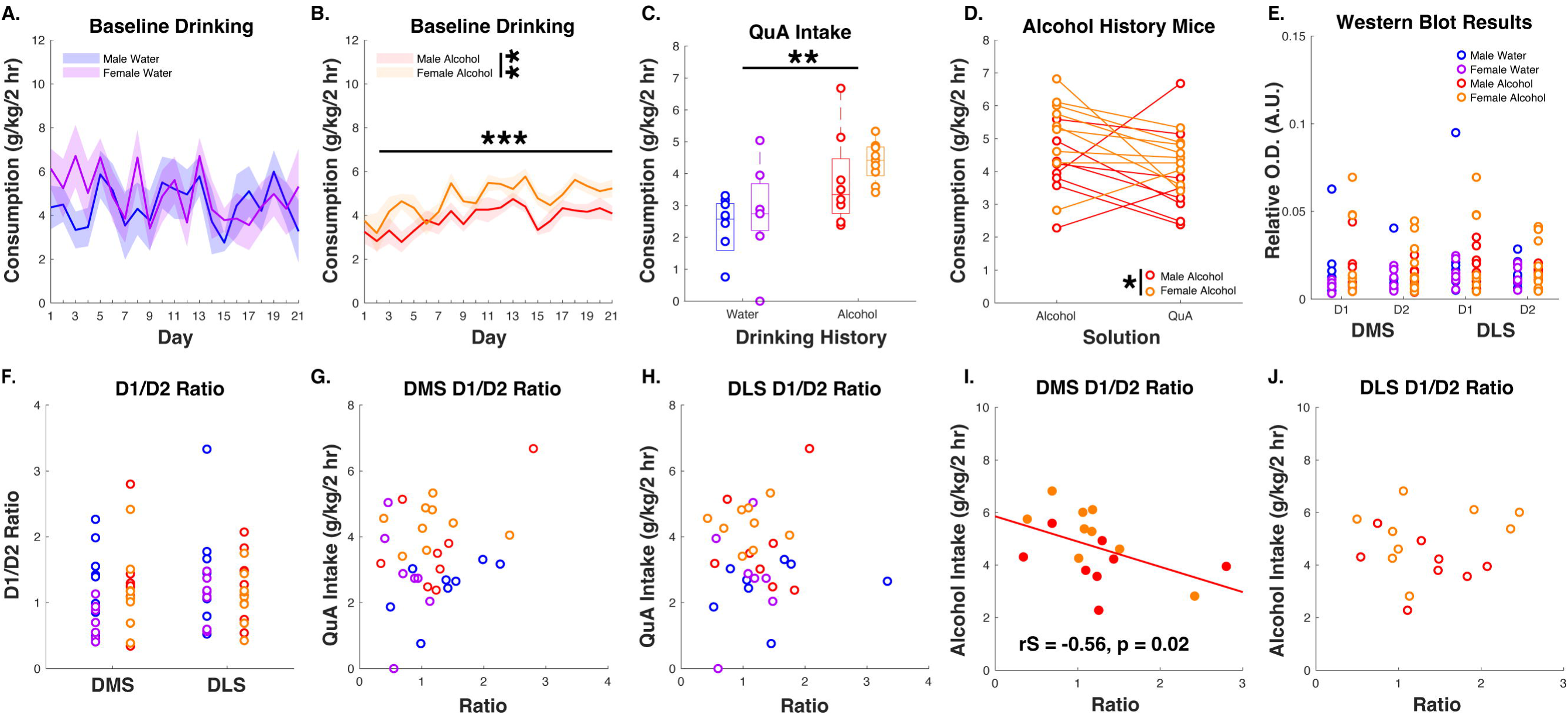
DMS D1/D2 Receptor Ratios Predict Alcohol, but not QuA, Consumption A-B. Male and female alcohol and water history mice drank for three weeks. On day 22 both groups were given QuA. **B.** RM ANOVA revealed a main effect of drinking history (***p < 0.05) and sex (** p < 0.05). **C.** Alcohol history mice drank more than water history mice (**p < 0.01). **D.** Alcohol history animals consumed the same amount of QuA as nonadulterated alcohol at baseline. Males drank less on average than females (*p < 0.05). **E.** D1 and D2 receptor expression in DMS and DLS tissue did not differ by sex or drinking history. **F.** D1/D2 ratios did not differ by sex or drinking history. **G-H.** QuA intake did not predict D1/D2 ratio for DMS or DLS. **I-J.** Alcohol intake predicts D1/D2 ratios in DMS (*p < 0.05), but not DLS. Data are displayed as individual data points or as mean ± SEM.

We next assessed the effect of an alcohol drinking history on ARD drinking (Figure 4C). RM two-way ANOVA of sex and drinking history on QuA drinking revealed a main effect of drinking history, F(1,28) = 12.28, p < 0.01, η²g = 0.3, and no significant effect of sex or interaction of sex and drinking history (p’s < 0.05). Next, we assessed the extent of the ARD in the alcohol history mice by comparing consumption of alcohol on the day prior to QuA access with QuA consumption. RM two-way ANOVA of sex and solution revealed a main effect of sex, F(1,15) = 4.57, p = 0.049, η²g = 0.15, Figure 4D.

D1 and D2 dopamine receptor expression in DMS and DLS tissue is shown in figure 1E. ANOVA of sex and drinking history for DMS D1, DMS D2, DLS D1, or DLS D2 (Figure 4E), and DMS D1/D2 ratios or DLS D1/D2 ratios (Figure 4F) revealed no significant effect (p’s > 0.05). Correlations of protein expression and QuA intake revealed no significant differences (data not shown). Correlations of DMS D1/D2 ratios (Figure 4G) and DLS D1/D2 ratios (Figure 4H) revealed no significant relationships with QuA intake. However, when we also assessed D1/D2 ratios and their relationship with alcohol drinking on the day prior to QuA intake, we found that DMS D1/D2 ratios had a strong negative relationship with alcohol intake in ARD mice, (r_S_ = -.56, n = 17, p = 0.02), Figure 4I. The DLS did not have a similar relationship (p > 0.05; Figure 4J). We also ran correlational analyses with water drinking on the day prior in the aversion-sensitive mice which revealed no differences and suggests that the above is not a general consummatory effect. These data suggest that the relative DMS D1 levels compared with D2 are related to alcohol, but not QuA drinking, in ARD mice and that lower D1/D2 ratios predict greater alcohol consumption in ARD mice.

## Discussion

The goal of this project was to elucidate whether the DMS and DLS engage in shifts in activity as a function of alcohol drinking history and the development of ARD. Based on previous research, we hypothesized that the DMS would disengage following the development of ARD and that the DLS would be more engaged following the development of ARD. In support of our hypothesis, we found a strong trend of DMS GluA1 AMPA receptors having a negative relationship with QuA drinking in aversion-sensitive mice. We hypothesized that this may be due to an increase in CP-AMPARs in the DMS which would act for rapid excitatory transmission which we thought acted as a “stop-signal” to prevent ARD. To causally test this, we microinjected NASPM, a CP-AMPAR antagonist, into the DMS in aversion-sensitive mice finding that intra-DMS NASPM significantly increased ARD, and that this effect was not due to a non-specific increase in consummatory behavior (water intake) or due to general alterations in DMS activity as M4 antagonists did not change QuA intake. We then hypothesized that control mice would have greater sensitivity to CP-AMPAR antagonists (NASPM) and show greater inward rectification than ARD mice. Counter to our hypothesis, we found that DMS SPNs of ARD mice were more sensitive to NASPM and had greater inward rectification than aversion-sensitive mice. Rectification index significantly predicted alcohol drinking history, but not QuA intake. Thus, our data suggest a complex and dichotomous relationship between CP-AMPARs in the DMS and ARD drinking. Finally, we found a negative relationship between DMS, but not DLS, D1/D2 ratios and alcohol drinking, but not QuA intake. Our findings provide evidence for a leading hypothesis stating that the activity in the DMS is required for preventing ARD. However, we also found evidence challenging the later portion of the hypothesis which posits a disengagement of the DMS once ARD has developed. Together, our results partially support our hypothesis and add complexity to the theories of addiction that suggest addiction is due to a shift away from DMS and towards DLS dependence finding both the DMS and DLS undergo changes that are related to ARD.

Since the DMS is necessary for flexible, goal-directed actions, DMS activity should act to drive goal-directed, adaptive, behaviors. In our experiment, we found that CP-AMPARs in the DMS act to prevent drinking despite negative consequences. When DMS CP-AMPARs were antagonized, aversion-sensitive mice drank QuA to intoxicating levels. This suggests Hebbian plasticity as a stimulus (QuA) results in rapid and functional alteration of AMPARs. Interestingly, when we recorded DMS SPNs, ARD mice had greater NASPM sensitivity (i.e. CP-AMPARs) compared with aversion-sensitive mice. This suggests two things: 1) increased CP-AMPARs on SPNs are a result of repeated binges, and 2) the CP-AMPAR activity necessary to prevent ARD might be localized on DMS cells other than SPNs. 5% of DMS neurons are interneurons and an unknown but potentially large proportion of all cells in the DMS are glial cells (von Bartheld et al. 2016). Glial cells are devoid of the GluA2 receptor and are therefore exclusively CP-AMPARS (Burnashev et al. 1992). Activation of glial AMPARs has been shown to cause vesicular glutamate release from the glial cells (Cervetto et al. 2015) and DMS astrocyte (a form of glia cell) activation alters DMS D1 and D2 SPN firing frequency (Kang et al. 2020). One potential hypothesis is that DMS glial cell activation is necessary for goal-directed behavior. Indeed, previous research has shown that activating DMS astrocytes restores goal-directed reward seeking in otherwise habitually responding animals (Kang et al. 2020). It is possible that the DMS CP-AMPAR activity gating ARD in our experiment is due to astrocyte activity and that as a result of repeated ethanol binges these astrocytes fail to do their basic function, clear extra glutamate, resulting in DMS SPNs undergoing plasticity to enhance SPN function via insertion of CP-AMPARs. This hypothesized mechanism could be caused by repeated ethanol access weakening astrocytic function while simultaneously enhancing SPN function which would account for our slice electrophysiology findings. However, it is unknown whether the DMS SPNs CP-AMPAR plasticity is the result of excitotoxity of SPNs or a gain of function via plasticity.

CP-AMPARs are both fundamental to central nervous system plasticity and function and can drive excitotoxicity, cell death, and diseases like addiction. Whether DMS CP-AMPARs are functioning in our experiments as a plasticity/gain of function or an excitotoxity mechanism resulting in ARD is unknown. Under the assumption the CP-AMPAR plasticity is driving greater synaptic weights, and thus, driving increased DMS reliance and function, DMS CP-AMPARs could be the site of increased motivation/goal-directedness resulting in the willingness of mice to continue to drink despite negative consequences. This hypothesis supports an alternative hypothesis of addiction which suggest that compulsive drug use is due to excessive goal-directedness (Hogarth, 2020) and that compulsive drug taking (as opposed to seeking) is due to enhanced motivation (Lüscher, Robbins, Everitt 2020). CP-AMPARs in the nucleus accumbens are the site of cocaine incubated craving (Conrad et al. 2008) where CP-AMPARS increase in number on SPNs in the NAc during cocaine withdrawal. When drug-paired cues are reintroduced resulting in glutamate influx to the NAc, the larger number of CP-AMPARs are thought to heighten the NAc’s response to glutamate, thus functionally driving craving and relapse following abstinence (Wolf 2016). A similar mechanism could be at play in the DMS where the DMS SPN CP-AMPAR plasticity following ARD could cause these neurons to become hyper-responsive to action-outcome and value information coming from cortical, thalamic, or substania nigra. We did not causally test this idea as we have previously had issues demonstrating robust ARD in cannulated mice (Bauer et al. 2022a). Future research using CP-AMPAR specific viral tools or cannulation in other behavioral models should investigate whether CP-AMPAR inhibition in ARD mice reduces ARD to provide further evidence surrounding the DMS’ role in ARD.

On the flip side of this coin, it is possible that the DMS loses normal function as a result of excitotoxicity via insertion of CP-AMPARs on DMS SPNs. This hypothesis fits with our finding that CP-AMPAR inhibition caused ARD in otherwise aversion-sensitive mice. Under this thinking, the DMS would lose normal functioning via repeated binges resulting in CP-AMPAR plasticity on SPNs that inhibit the ability of the DMS to rapidly modulate goal-directed behavior via excitotoxicity of DMS synapses. Previous research has shown that systemic administration of AMPAR antagonists (Stephens and Brown, 1999; Ruda-Kucerova et al. 2018; Bauer et al. 2020) can reduce alcohol drinking which provides support that ethanol exerts some of its effects through glutamate induced excitotoxicity, which has long been considered (Lovinger 1993). However, we did not explicitly measure biomarkers of excitotoxity or cell death within the DMS. To our knowledge, there has been no assessment of the relationship between excitotoxicity and ARD.

The DMS is composed of a variety of neurons including interneurons (cholinergic, fast spiking, and GABAergic) as well as primarily SPNs. The SPN are categorized based on whether they express primarily D1 or D2 areceptors and their projection pathway to the substantia nigra reticula (SNr)/globous pallidus internus (GPi) and globous pallidus externus (GPe), respectively (Gerfen 1990). D1 SPNs, D2 SPNs, and interneurons both work together to coordinate behavior as well has have leading roles in specific behaviors. The results of our experiments support the idea that each receptor and cell types have leading roles in specific behaviors. For example, we found that DMS CP-AMPAR, but not DMS M4 receptor antagonism altered QuA intake. M4 antagonism reduced locomotor activity without altering QuA consumption. We also found that DMS D1/D2 ratios negatively predicted alcohol, but not QuA, consumption in ARD mice. The DMS D1/D2 ratio did not predict water consumption in aversion-sensitive mice. We did not assess D1 vs D2 SPN in our electrophysiology recordings. However, given that our electrophysiology results have very little heterogeneity, it is plausible that D1 vs D2 cells do not differentially respond to the CP-AMPAR manipulations. Interestingly, others have found D1 DMS SPNs to be drivers of ARD (Houck 2019; da Silva et al. 2024). Our findings likely demonstrate that general DA ratios may modify as a function of repeated binges but that a more precise targeting (like with optogenetics or specific DA antagonist) may be necessary to identify functional roles of D1 and D2 neurons in the DMS for ARD. Together, these data demonstrate specific roles for DMS M4, CP-AMPARs, and D1/D2 ratios on behavior.

Finally, the overarching purpose of this project was to identify differences between the DMS and DLS over the development of ARD. We found that the DMS is functionally involved in ARD and that the DMS undergoes plasticity changes on the SPNs as a function of alcohol experience. We were intrigued by the negative correlation between GluA1 and QuA consumption in aversion-sensitive mice which propelled us to investigate the DMS and not both the DMS and DLS. However, it is worth noting that female ARD mice had a significant positive relationship between QuA and GluA1/A2 suggesting that the DLS also undergoes some sort of plasticity in association with ARD. The overall scope of this paper is limited by the lack of causal assessment of the DLS in ARD. However, research from our lab has shown that alcohol binge-drinking is initially sensitive to intra-DLS AMPAR antagonism, but that sensitivity is gone following extended binge drinking episodes that result in ARD or innate ARD (Bauer et al., 2021; 2022a; 2024). Others have found changes in DLS plasticity following alcohol exposure (Corbit, Nie, Janak 2012; 2014; Rangel-Barajas 2021; Jeanblanc et al. 2009; DePoy et al. 201; Muñoz et al. 2018; Haggerty et al. 2022) and the DLS is necessary for ARD (Giuliano et al. 2019; Patton et al. 2021, Bauer et al., 2021]. We note that we provide some evidence for DLS involvement in ARD despite a primary focus on the DMS, the DLS is also likely involved in ARD, though whether CP-AMPARs are the basis of DLS-dependent ARD is unknown.

In conclusion, we have identified a pharmacological underpinning of ARD within the DMS. CP-AMPARs act to prevent ARD in a specific manner, however SPNs, the primary cell type within the DMS, do not innately express CP-AMPARs, which suggests a different neuron or cell type is acting to prevent ARD. After repeated binges that result in ARD, SPNs in the DMS express CP-AMPARs and have inward rectification. The inward rectification of the SPNs does not predict ARD but does predict baseline alcohol drinking with could suggest that the SPN modification are not related to ARD directly. Finally, we found that the DLS may also undergo plasticity relating to ARD though we did not explicitly explore cell-dynamics within the DLS. Lastly, these data provide evidence that the DMS is involved in goal-directed action prior to the development of ARD and that the DMS undergoes plasticity that could be interpreted as a gain of function following the development of ARD which provides support for excessive motivation and goal-directedness during ARD. Ultimately, we found no evidence that the DMS disengages during ARD though future research should causally test such hypotheses.

## Author Contributions

Conceptualization: Meredith Bauer, Claudia Rangel-Barajas, Stephen Boehm

Methodology: Meredith Bauer, Claudia Rangel-Barajas, Yanping Zhang, Stephen Boehm

Formal Analyses and Investigation: Meredith Bauer, Claudia Rangel-Barajas, Yanping Zhang

Writing - original draft preparation: Meredith Bauer

Writing – Review and editing: Claudia Rangel-Barajas, Yanping Zhang, Stephen Boehm

Funding acquisition: Stephen Boehm

## Data Availability Statement

The data are freely available upon reasonable request to the corresponding author

